# Transcription-coupled repair and mismatch repair contribute towards preserving genome integrity at mononucleotide repeat tracts

**DOI:** 10.1101/584342

**Authors:** Ilias Georgakopoulos-Soares, Gene Koh, Josef Jiricny, Martin Hemberg, Serena Nik-Zainal

## Abstract

The mechanisms that underpin how insertions or deletions (indels) become fixed in DNA have primarily been ascribed to replication-related and/or double-strand break (DSB)-related processes. We introduce a novel way to evaluate indels, orientating them relative to gene transcription. In so doing, we reveal a number of surprising findings: First, there is a transcriptional strand asymmetry in the distribution of mononucleotide repeat tracts in the reference human genome. Second, there is a strong transcriptional strand asymmetry of indels across 2,575 whole genome sequenced human cancers. We suggest that this is due to the activity of transcription-coupled nucleotide excision repair (TC-NER). Furthermore, TC-NER interacts with mismatch repair (MMR) under physiological conditions to produce strand bias. Finally, we show how insertions and deletions differ in their dependencies on these repair pathways. Our novel analytical approach reveals new insights into the contribution of DNA repair towards indel mutagenesis in human cells.

## Main text

While substitutions in human cancers have been extensively studied, insertions/deletions (indels) have remained comparatively under-explored. This was historically due to the relative difficulty in obtaining high-quality indel data, further restricted by a limited repertoire of approaches to analyse indels as extensively as substitutions. Nevertheless, indels are common in human cancers and their location and sequence composition are non-random. Thus, like substitutions, they provide important insights into the mutational processes that have shaped the landscape of cancer genomes.

## Landscape of insertions and deletions across human cancers

We utilized 2,416,257 indels from a highly curated set of 2,575 whole-genome sequenced (WGS) cancers of 21 different cancer-types. Median indel number per tumour was 386, corresponding to a conservative indel density of 0.127 per Mb per cancer genome. Deletions (median 222) were more prevalent than insertions (median 124) in the majority of cancers (Figure 1a, Supplementary Figure 1a, ratio of deletions to insertions per tumour ranges from 0.8 to 4.39). Moreover, deletion size showed greater variability than insertion size across and within tumour-types (Figure 1b-c, Supplementary Figure 1b).

**Figure 1:**
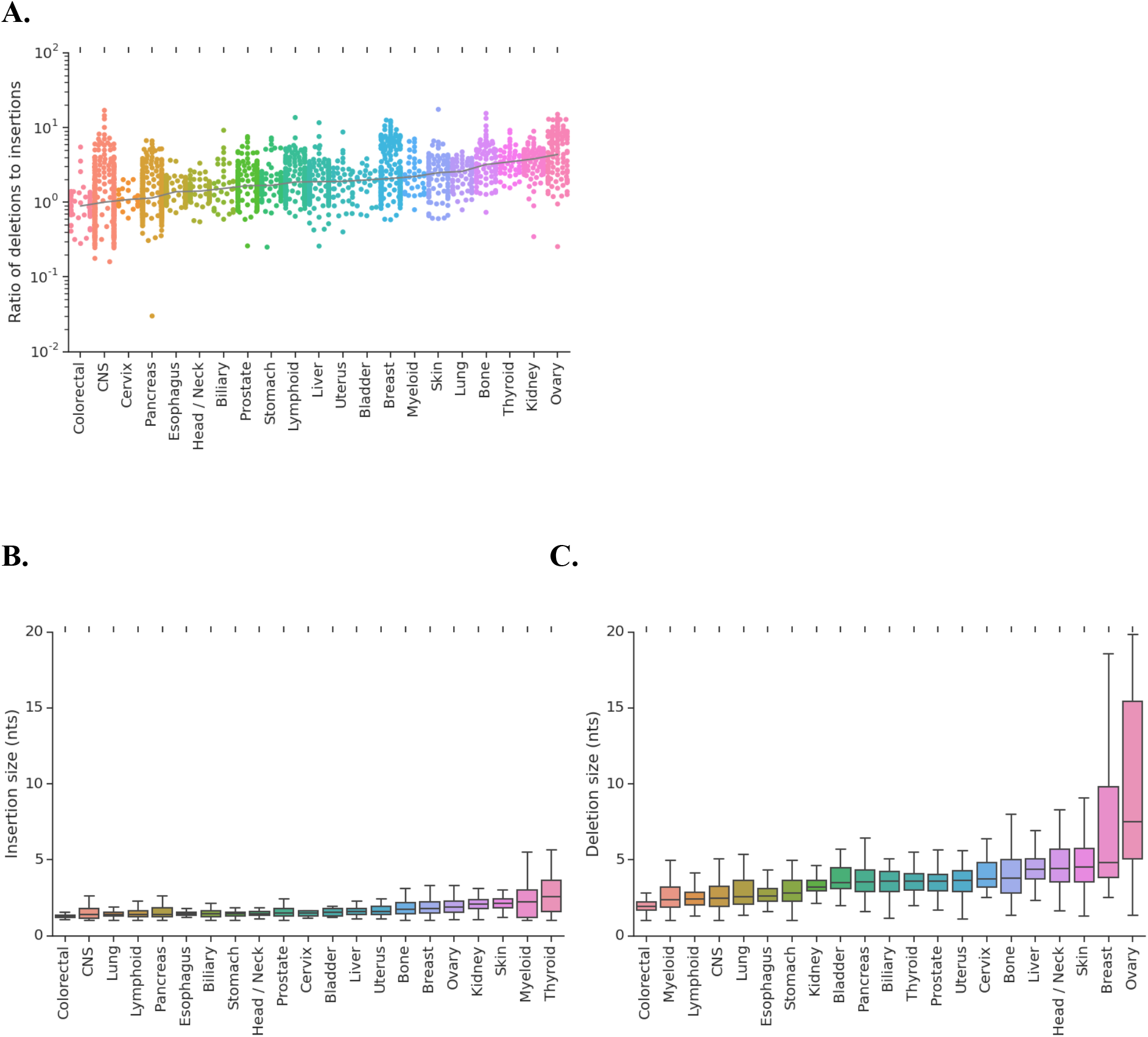
Indel characteristics across cancer types. **A)** The ratio of deletions to insertions for each tumour-type. **B)** Distribution of size of insertions for each tumour-type. **C)** Distribution of size of deletions for each tumour-type.

This first observation can be broadly explained by already-known mechanisms that generate indel lesions. Replication-related DNA polymerase slippage errors running through microsatellites tend to cause deletions (Strand et al. 1995), because single-stranded DNA ahead of a polymerase can twist, causing a single repeat unit of a run of mono- or dinucleotides in the template strand to loop out. A polymerase passing over such a loop would generate a deletion (Strand et al. 1995), (Tran et al. 1996), (Tran et al. 1997), (Sia et al. 1997). Because these small insertion/deletion loops (IDLs) are efficiently repaired by MMR, the density of deletions in microsatellites in higher in cells lacking MMR. This phenomenon is referred to as microsatellite instability (MSI). The likelihood of formation of such loops increases with the length of the repeat and we confirm this by showing that indel frequency increases with increasing lengths of polynucleotide repeat tracts (Supplementary Figure 1c-d) and is more pronounced in MMR-deficient samples. Double-strand breaks (DSB) can give rise to deletions if repaired by non-homologous end-joining (NHEJ), or if addressed by homology-directed sub-pathways such as single-strand annealing (SSA) or microhomology-directed end-joining (MMEJ) (Delacôte et al. 2002), (Simsek and Jasin 2010), (Nik-Zainal et al. 2012), (Jasin and Rothstein 2013), (Ghezraoui et al. 2014), (Kass et al. 2016), (Nik-Zainal et al. 2016). The latter result in larger deletions (3bp in size or more), thus explaining the broader spectrum of observed deletion sizes (Figure 1c).

In contrast, a transient dissociation of the primer and template strands and reannealing of the primer in a wrong register within the microsatellite could cause both an insertion or a deletion, but these are less likely to arise during normal replication (Petrov 2002), because the end of the primer strand is tightly bound by the replisome. Thus, the occurrence and the relative frequencies of indels and their size spectra can be explained by known mechanisms.

## Transcriptional strand asymmetry of indels

In addition to classical contributions of replication and DSB-repair pathways to indel formation, we introduce another dimension to exploratory analyses of indel mutagenesis: the contribution of transcription. Transcription has been implicated in asymmetric distribution of substitutions between strands for decades (Francino et al. 1996), (Majewski et al. 2003), (Morganella et al. 2016). In particular, transcription-coupled nucleotide excision repair (TC-NER) is believed to preferentially repair DNA damage on the template (transcribed or non-coding) strand. TC-NER activity is thus inferred from the excess of mutagenesis on the non-template (coding) as compared to the template (non-coding) strand, particularly for those environmental mutagens where the target of primary DNA adduct formation is known. For example, guanines adducted by tobacco carcinogens result in an excess of G>T mutations on the non-template strand (Denissenko et al. 1998), (Rodin and Rodin 2005), (Pleasance et al. 2010). Likewise, primary covalent modifications of cytosines forming 6,4 pyrimidine-pyrimidone dimers (6,4-PPs) and cyclobutane pyrimidine dimers (CPDs) by ultraviolet light are preferentially repaired on the template strand resulting in an excess of C>T transitions on the non-template strand (Fousteri and Mullenders 2008). However, to the best of our knowledge, transcriptional strand asymmetry in indels has not been investigated, primarily due to the technical challenge of being unable to orientate each indel with respect to transcriptional strand.

We set out to determine transcriptional strand asymmetry of indels by focusing on mononucleotide repeats of up to ten base pairs in length. We first analysed the distribution of mononucleotide repeats across the gene body in the reference human genome. Each gene was divided into ten equal-sized bins to correct for differences in gene length. Two additional bins were added upstream of the transcription start site (TSS) and two downstream of the transcription end site (TES), each 10kB in length, resulting in a total of 14 bins.

### Asymmetries of repetitive tracts in the reference genome

We observed a strong enrichment of polyG/polyC motifs directly upstream and downstream of the TSS and downstream of the TES; this contrasted with the distribution of polyT/polyA motifs, which were found to be enriched throughout gene body (Figure 2a).

**Figure 2:**
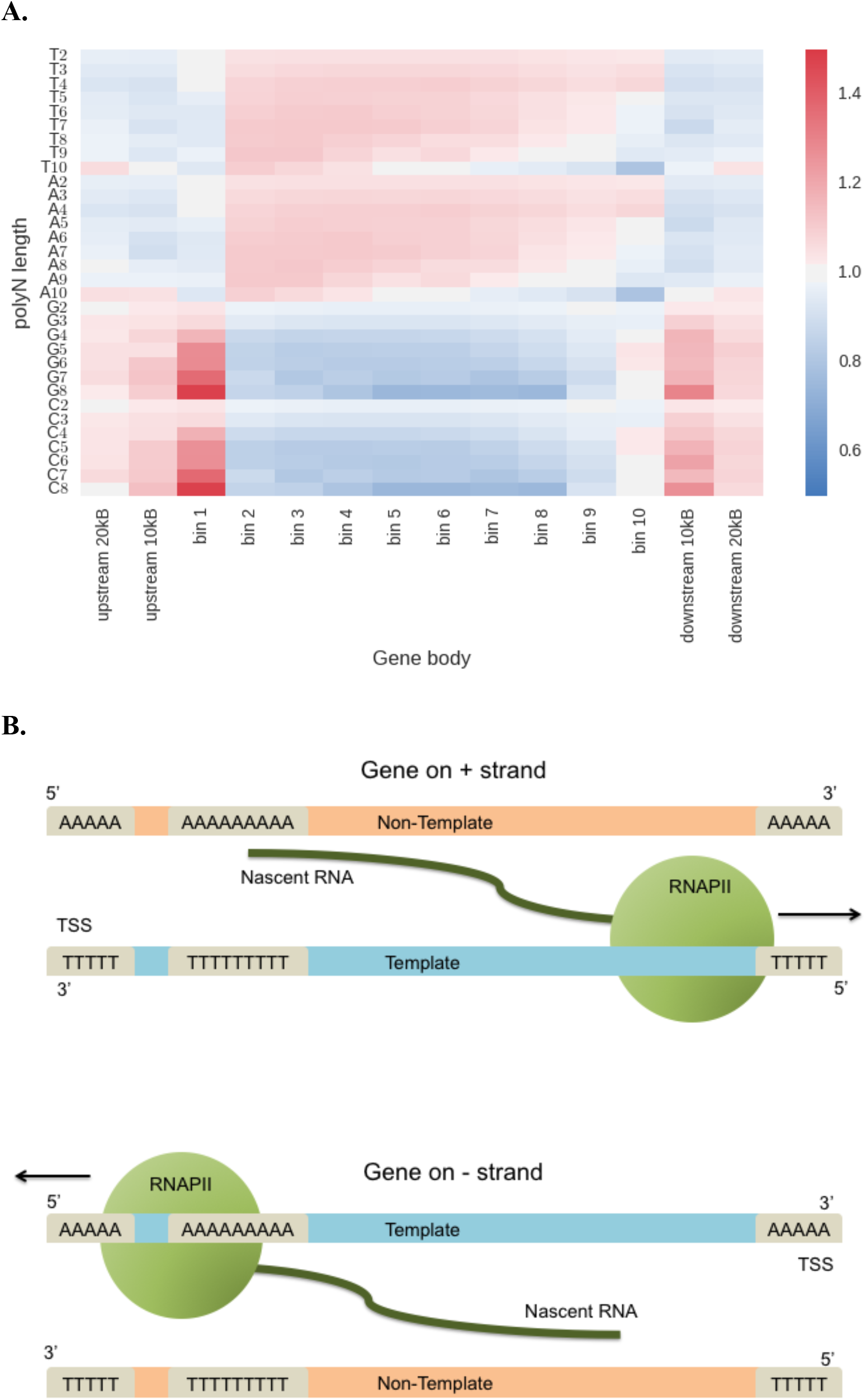
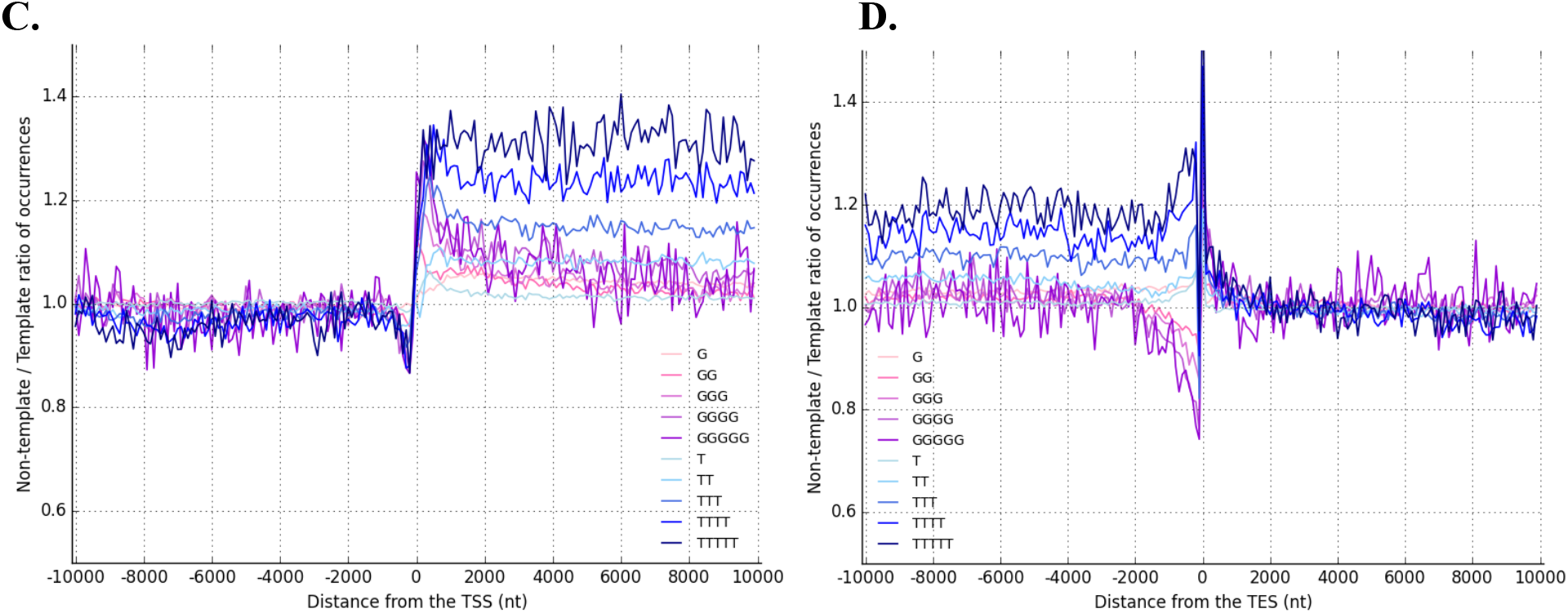
Strand asymmetries of polynucleotide (polyN) repeat tracts within transcribed regions. **A)** Density of various polyN motifs across genes. Each gene is divided into ten bins, and two additional bins are added at either end of each gene. For any given bin, red indicates relative enrichment in comparison to all other bins for that polyN, whereas blue indicates relative depletion. **B)** Scheme depicting the identification of polyN motifs on the template (blue) or non-template (orange) strands, dependent on the direction of the gene. RNA-polymerase II (RNAPII) binds to the template strand and mediates transcription. Thus, in the panel above, where the gene is on the (+) strand, the polyA tracts are on the non-template strand. In the panel below, where the gene is on the (–) strand, the polyA tracts on the template strand. **C)** Density of polyT and polyG motifs around the transcription start site (TSS). The gradient of pink to purple represents polyG tracts of 1-5bp length, whereas the gradient of light blue to dark blue represents polyT tracts of 1-5bp length. **D)** Density of polyT and polyG motifs around the transcription end site (TES).

We calculated the frequencies of each polyN motif (where N is any nucleotide) on the template and non-template strands in the reference genome. Because the direction of transcription for each gene in the genome is known, each polyN motif can be orientated (Figure 2b). For example, for a gene on the (+) strand, the template strand is the (-) strand. A polyT motif that is on the (-) strand of this gene is therefore described as being on the template strand. It can also be described as a poly-A motif on the non-template strand (Figure 2b). Using this reasoning, we assigned each polyN motif to either the template or non-template strand of the reference genome. If there were no asymmetries, polyN tracts would occur with equal probabilities on either strand.

Intriguingly, we found that polyT motifs displayed a bias towards the non-template strand in the reference genome (Figure 2c-d), with a non-template to template asymmetry enrichment for short polyT motifs of ~1.15-fold. This was tract-length-dependent, where longer repetitive tracts were associated with greater strand bias (Figure 2b, Supplementary Figure 3a-e) up to ~1.4-fold at >5nt polyT motifs (weighted average asymmetry of 1.14-fold, Supplementary Figure 3a). In contrast, we did not observe a similarly pronounced asymmetry of polyG motifs across gene bodies, although a skew in polyG motifs was noted at the boundaries of gene bodies, fitting with previous reports of a high GC-skew at either end of genes (Ginno et al. 2013) (Figure 2c-d, Supplementary Figure 2a-b, Supplementary Figure 3c-e). The marked variation in strand distributions of the polyN motifs in the reference genome can be visually appreciated particularly around the TSS and TES (Figure 2c-d, Supplementary Figure 3c-e).

### Transcriptional strand asymmetries of small indels occur at polynucleotide repeat tracts

We next investigated whether there was strand asymmetry in the occurrence of indels at these polyN tracts, correcting for the skewed background distributions of polyT and polyG motifs. Across cancers, polyT motifs of 2-10bp in length were consistently more mutable on the nontemplate strand, but the levels of asymmetry varied by cancer type, with an increase in the likelihood of indel mutagenesis on the non-template over the template strand ranging from 2.1% in ovarian cancers to 16.5% in uterine cancers (Figure 3a, e). This finding was surprising, given that the prevailing dogma on indel formation – particularly at poly-nucleotide repeat tracts – involves the formation of small IDLs that are substrates for MMR (Ellegren 2004), (Garcia-Diaz and Kunkel 2006), (Romanova and Crouse 2013), (Kunkel and Erie 2015). However, our analysis showing marked transcriptional strand asymmetry implies either the activity of TC-NER at these motifs or the activity of transcription-associated damage.

**Figure 3:**
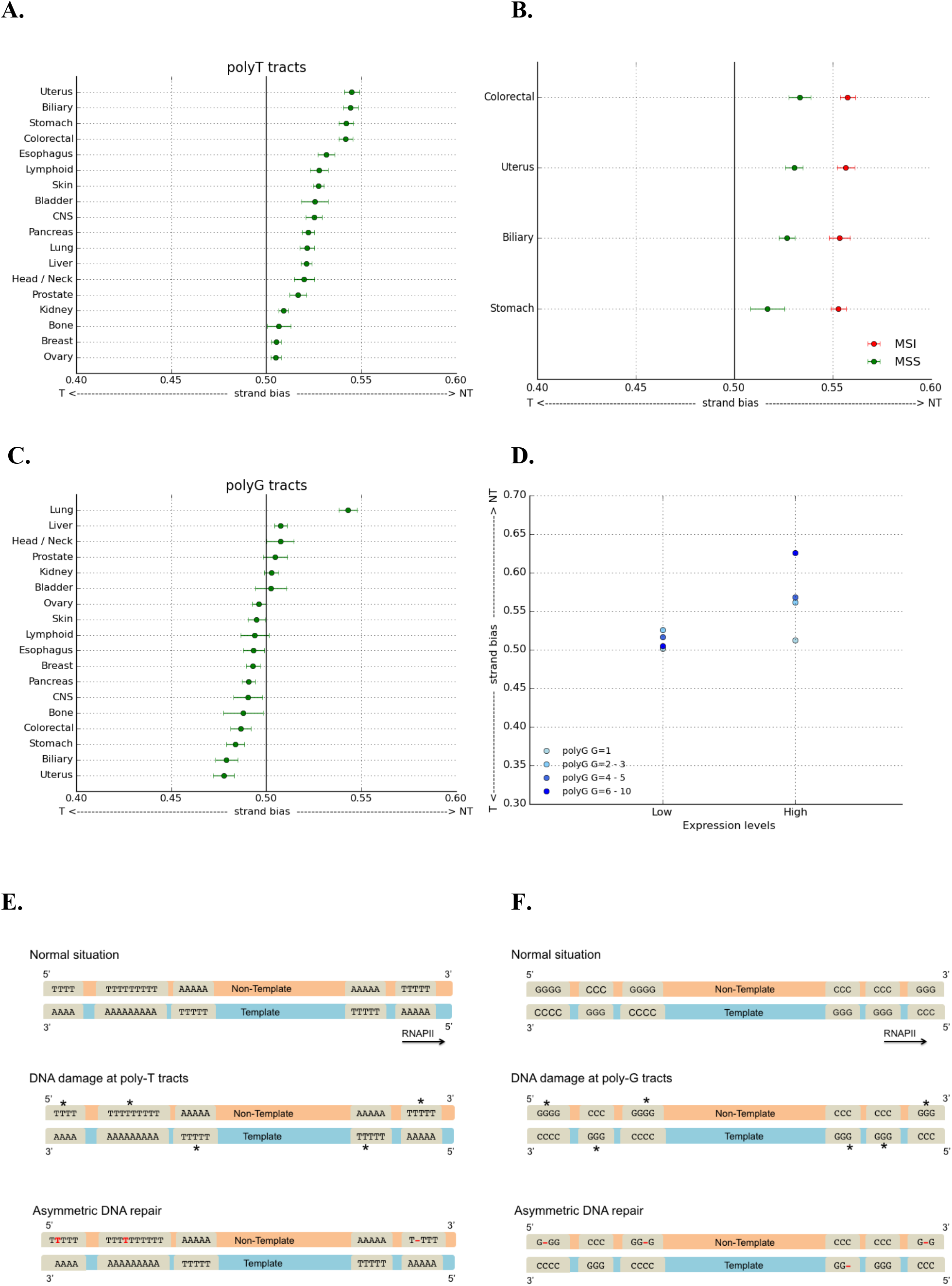
Transcriptional strand asymmetry of indels that occur at polyN tracts across multiple cancer types. **A)** Bias towards the non-template strand at polyT motifs for indels. Average bias is shown with error bars showing standard deviation from 1,000 bootstraps. Myeloid, cervix and thyroid cancers were excluded due to low numbers of total indels (Supplementary table 1). **B)** MSI samples display stronger non-template strand bias for stomach, biliary, uterus and colorectal tumours. **C)** Transcriptional strand asymmetries of indels at polyG motifs. Average bias is shown, with error bars showing standard deviation from bootstrapping. **D)** Non-template strand bias for indel formation at polyG motifs in lung cancer is associated with gene expression levels and length of polyG tracts. **E)** Scheme depicting mechanism of indel mutagenesis at poly-T tracts. DNA damage, shown as asterisks (*) that arise at T nucleotides of poly T tracts can occur on both template and non-template strands. The subsequent DNA repair, postulated to be TC-NER, results in preferential correction of DNA damage on the template strand, leaving T insertions (highlighted in as red T’s) and T deletions (shown as red -) on the non-template strand. **F)** Schematic depicting mechanism of indel mutagenesis at poly-G tracts in lung cancers from smokers. DNA damage in the form of adducted guanines (*) are asymmetrically repaired by TC-NER, with preferential repair of the template strand, thus accumulating more G indels on the non-template strand.

We noted that uterine, colorectal, biliary and stomach cancers showed the highest levels of transcriptional strand asymmetry, with a 16.5%, 15.5%, 16.3% and 15.5% higher frequency of indels occurring at polyT motifs on the non-template strand compared to the template strand, after correcting for background differences in distribution of polyT motifs (Figure 3a). These cancer types are often associated with a high frequency of MMR deficiency (Baretti et al. 2018). To explore the contribution of MMR to transcriptional strand asymmetries of indels at polyT tracts, we compared samples with MMR deficiency and MSI to microsatellite stable (MSS) samples. Surprisingly, in MSI samples, transcriptional strand bias for polyT motifs towards the nontemplate strand was more pronounced in comparison to MSS samples, with a 7.9-12.8% higher frequency of indels (Figure 3b). This suggests that not only is TC-NER implicated in the repair at polyN motifs, it is also dependent on the normal physiological functioning of MMR. In the absence of MMR, damage at these sites relies more heavily on TC-NER alone, resulting in an increase in strand bias.

To validate this hypothesis regarding the reliance of TC-NER on MMR, we examined experimentally-generated mutation patterns from CRISPR-Cas9 knockouts of a human cancer cell line, HAP1 (Zou et al. 2018). We would expect the presence of transcriptional strand bias under normal conditions but that the magnitude of the effect would be increased in MMR gene knockouts. Indeed, our analytical findings are recapitulated in the experimental setting. In a knockout model of MutS homolog 6 *(MSH6)*, a key MMR gene, in total 1,663 indels occurred on the non-template strand polyT tracts, whereas 1,165 indels occurred on template polyT tracts, corresponding to a 16.9% corrected increase in frequency on the non-template strand (p-value<0.001), a similar magnitude to that observed in cancers.

TC-NER was not previously thought to play a role in the maintenance of genome integrity at polynucleotide repeat tracts. Interestingly, we did not observe strong transcriptional strand bias for indels overlapping polyG motifs of 2-10bp in length. Lung cancers were a notable exception; they exhibited a large excess of G indels on the non-template strand (15.8% higher frequency of indels at polyGs on the non-template strand when compared to those on the template strand) (Figure 3 c). This pattern of asymmetry mirrors what was observed for G>T substitutions in lung cancers, which are attributed to the formation of bulky adducts on guanines from tobacco-related carcinogens. This type of helix-distorting damage is classically repaired by TC-NER (Denissenko et al. 1998), (Rodin and Rodin 2005), (Pleasance et al. 2010). The observation of transcriptional strand asymmetry for G indels at polyG tracts in tobacco-associated lung cancers reinforces the involvement of TC-NER in the metabolism of polyN motifs.

To validate this observation, we analysed indel mutational profiles of non-cancerous human cells exposed to various polycyclic aromatic hydrocarbons (PAHs) including benzo[a]pyrene [0.39 μM and 2 μM] and benzo[a]pyrene diol epoxide [0.125 μM], believed to be the carcinogenic components of tobacco smoke. We observed that 77 indels occurred on non-template polyG tracts, in contrast to only 39 indels on template polyG tracts. This corresponds to a 94.1% increase in the frequency of indels at the non-template over the template strand, corrected relative to the expected frequency and based on the background distribution of polyG tracts. This supports our analytical observations of *in vivo* patterns derived from studying human cancers (Figure 3c, f).

The activity of TC-NER is linked to gene expression levels (Hanawalt and Spivak 2008) where higher levels of transcription are associated with increased TC-NER activity. To further support our findings that TC-NER plays a role in the repair of polyG motifs in lung cancers, we explored the degree of asymmetry in relation to gene expression levels. We used gene expression data from a representative cell-of-origin (Supplementary Table 2). In keeping with our hypothesis that TC-NER plays a pivotal role in the repair at polyG tracts, there was minimal transcriptional strand asymmetry for polyG motifs at genes that were not expressed or lowly expressed, and strong asymmetry for highly expressed genes. This was also strongly dependent on the length of the polyG motifs (Figure 3d).

### Insertions and deletions are differentially-dependent on DNA repair pathways

Next, we distinguished insertions from deletions at polyT and polyG tracts to find that transcriptional strand asymmetry differed between these classes of indels (Figure 4a, Supplementary Figure 4a-b). Insertions showed aggravated asymmetries at polyT tracts across all cancer types (Figure 4a), suggesting that mutagenesis associated with polyT tracts is highly dependent on TC-NER. By contrast, non-template strand bias of deletions at poly-T tracts was restricted to tumour types that had a high incidence of MSI (biliary, colorectal, stomach and endometrial). The levels of asymmetry for insertions was independent of MSI (Figure 4a). Thus, mutagenesis that results in deletions is more heavily dependent on the MMR pathway.

**Figure 4:**
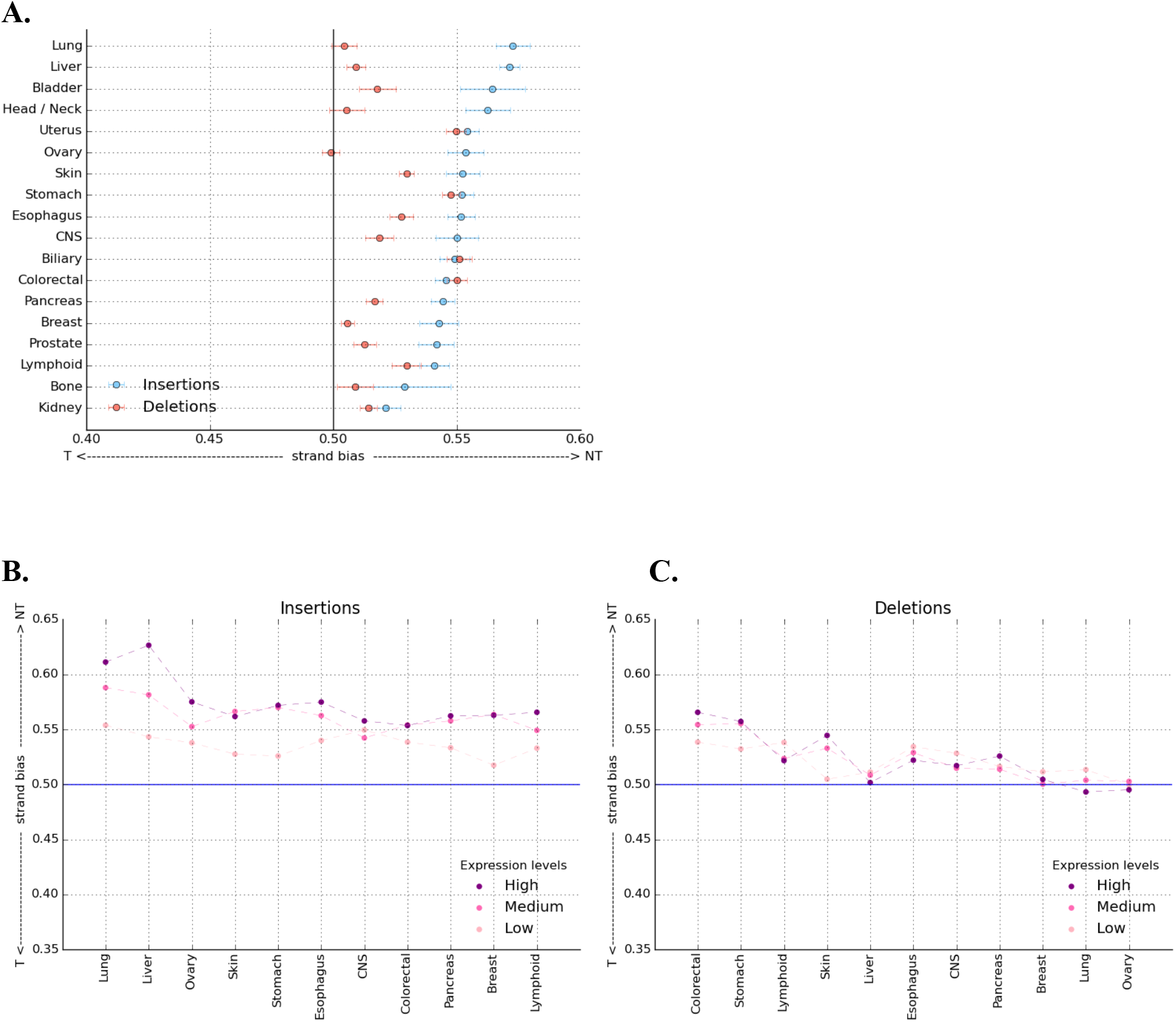
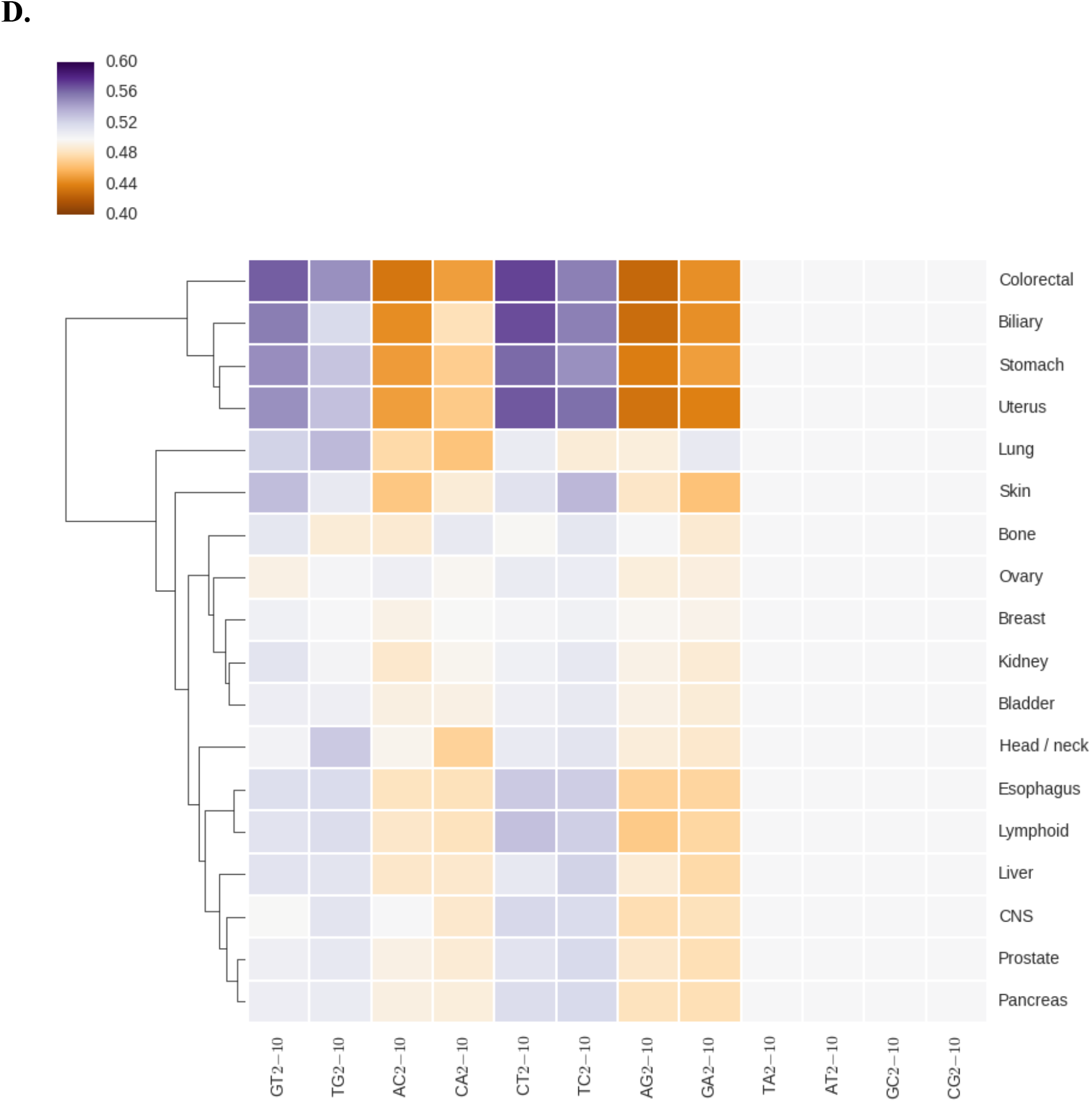
Transcriptional strand asymmetry at insertions and deletions at transcribed regions. **A)** Transcriptional strand asymmetry of insertions and deletions at polyT tracts. Error bars represent standard deviation from bootstrapping with replacement. **B)** Transcriptional strand asymmetry occurring at polyT tracts according to level of gene expression for insertions and **C)** deletions across tumour types. **D)** Hierarchical clustering displaying transcriptional strand asymmetries for indels overlapping dinucleotide motifs. Dinucleotide repeat tracts of up to five repeated units are displayed. Purple represents asymmetry towards the non-template strand, whereas orange represents asymmetry towards the template strand. In the dendrogram of cancers, biliary, uterus, colorectal and stomach cancers are more distant from the other cancers, and contain MSI samples, while lung cancers are also separable from other cancer types, further reinforcing our observations regarding the DNA damage and repair processes that contribute to the observed asymmetries.

To support this hypothesis, we investigated the relationship between transcriptional strand asymmetry for insertions and deletions at polyT motifs, and gene expression levels (Supplementary Table 2). Genes with higher expression levels displayed stronger transcriptional strand asymmetry of insertions at polyT tracts across all inspected cancer types (Figure 4b), implicating TC-NER, which is linked to expression levels (Hanawalt and Spivak 2008). However, a relationship between expression levels and transcriptional strand asymmetry of deletions at polyT tracts could only be observed for a subset of cancer types (Figure 4c) and the strand bias was less apparent. In contrast, at polyG motifs, we did not observe consistent associations between expression levels and transcription strand asymmetry of insertions or deletions across cancer types (Supplementary Figure 4c-d), with the exception of lung cancers; this we expected because of the influence of mutagenesis linked to tobacco carcinogens (Figure 3d, Supplementary Figure 4c-d).

### Transcriptional strand asymmetries at indels may be a general feature

Finally, to understand whether our observations were restricted to mononucleotide tracts or if they could be a more generic mechanistic feature of indel mutagenesis, we attempted to explore other types of indels. The limitation is the difficulty to assign other categories of indels or genomic characteristic to specific stands. It was, however, possible to ascribe strandedness to dinucleotide repeat tracts. There were some caveats: palindromic GC/CG and AT/TA dinucleotides could not be oriented, and AA/TT/GG/CC dinucleotides were excluded because these are similar to mononucleotide polyA/T/G/C’s respectively. This left us with eight types of poly-dinucleotide repeat tracts that we could analyse (GT/TG/AC/CA/CT/TC/AG/GA), (Supplementary Figure 5). Indeed, correcting for background asymmetries in the genome, we observed aggravated transcriptional strand asymmetries for several poly-dinucleotide repeat tracts (Figure 4d). This was most marked amongst tumour types where MSI was prevalent (Figure 4d, Supplementary Figure 6). Furthermore, strand asymmetry in insertions was stronger than in deletions, but the opposite was true for MSI tumours (Supplementary Figure 7a-b). Thus, our findings appear to be applicable to motifs other than mononucleotide repeat tracts.

## Conclusion

In this work, we have described a novel method to investigate transcriptional strand asymmetries for indels. Unexpectedly, we found biases in the distribution of mononucleotide repeat tracts in the reference genome at transcribed regions, and the bias is more pronounced for longer tracts. This bias needs to be considered when exploring transcriptional strand asymmetries for indels overlapping mononucleotide repeat tracts.

Our analysis demonstrates strong and previously-undescribed transcriptional strand asymmetries of indels. Our results implicate particular DNA repair pathways, namely TC-NER and MMR as factors contributing factors to the observed strand biases at indels (Figures 3–4). We further reveal that the formation of insertions is entirely TC-NER-dependent, while the formation of deletions is additionally reliant on MMR, thus reinforcing how distinct mechanisms underpin the formation of different classes of indel.

## Materials and Methods

### Mutation calling

Data were obtained from whole genome sequenced (WGS) cancers from ICGC under the project Pan Cancer Analysis of Whole Genomes (PCAWG). They included 46 cancer projects from 21 organs. In total, 2,575 whole genome sequenced patients were analysed using the GRCh37 (hg19) reference assembly of the human genome.

Somatic indel calls were performed using three pipelines from four somatic variant callers. These were the Wellcome Sanger Institute pipeline, the DKFZ/ EMBL pipeline and the Broad Institute pipeline. Indels were identified as described in (Campbell et al. 2017), with somatic variant false discovery rate of 2.5%. Indel calling was performed by those algorithms and only indels called by at least two of the callers were analysed (Campbell et al. 2017), therefore generating a conservative dataset (Supplementary Table 1). As a result, the false negative rate of indel detection could be higher than that of other methods, and of each pipeline separately, which implies that many indels present in the samples were not identified successfully. However, because of the large number of WGS tumour samples available, a sufficient number of indels remained (Supplementary Table 1). Finally, for a small subset of indels, the indel calls were visually examined using JBrowse Genome Browser (Buels et al. 2016), to inspect the number of reads reporting the indel, if the indel calls were biased towards the end of the sequencing reads or if there were other systematic biases between the normal and tumour sequencing reads; such biases could not be identified.

The distance between each pair of consecutive indels was calculated per patient. Indels in different chromosomes were excluded since we could not define their pairwise distance. Patients without multiple indels in the same chromosome were excluded. The same analysis was also performed separately for insertions and deletions to generate Supplementary Figure 1a.

Indels from HAP1 cells with MutS homolog 6 *(MSH6)* knock-out were obtained from (Zou et al. 2018). Indels from cells exposed to various polycyclic aromatic hydrocarbons (PAHs), namely benzo[a]pyrene [0.39 μM and 2 μM] and benzo[a]pyrene diolepoxide [0.125 uM], were obtained from (Zou et al. in press) to examine transcriptional strand asymmetry at indels overlapping polyN motifs in experimental settings.

Substitution calling was performed as described in (Campbell et al. 2017). For lung cancers, C>A substitutions were examined with respect to transcriptional strand asymmetries at polyG tracts and replication timing (Supplementary Figure 4e).

Mutational enrichment at MSI over MSS samples was defined as: Ratio: (Proportion of indels overlapping polyA/T at MSI samples) / (Proportion of indels overlapping polyA/T at MSS samples)

### Template / Non-template strand asymmetries at the reference human genome

Gene annotation from Ensembl was followed (Aken et al. 2016). BEDTools utilities v2.21.0 were used to manipulate genomic files and intervals (Quinlan and Hall 2010). GC-skew is a measure of bias in the number of Gs or Cs between the template and non-template strands. GC-skew was calculated as (G-C) / (G+C) for windows of 100 bp around the TSS and TES. Similarly, AT-skew was calculated as (A-T) / (A+T) for windows of 100 bp around the TSS and TES (Supplementary Figure 2a-b).

Genes in the positive and negative orientations were separated to determine the direction of gene transcription. Scripts were written in python to identify non-overlapping polyN motifs of size 1-10bp as well as dinucleotide motifs of length 2-10bp genome-wide and orient them in terms of transcription direction at genic regions.

Template motifs were the motifs in: i) positive gene orientation and negative genome strand, ii) negative gene orientation and positive (reference) genome strand. Non-template motifs were the motifs in: i) positive gene orientation and positive genome strand (reference), ii) negative gene orientation and negative genome strand. Bedtools *intersect* utility was used to calculate motif occurrences in template and non-template strands across genic regions.

To investigate the effect of the distance from the TSS and the TES across the gene length, for genes with unequal gene length, we divided each gene into ten genomic bins of equal size. Also, two additional bins upstream from the TSS and two bins downstream of the TES, each 10kB in size, were added. Then, we calculated the frequency and the strand asymmetry bias of polyN motifs in each genic bin (Figure 2a, Supplementary Figure 3).

Relative enrichment of a polyN tract at a bin was calculated as:

Enrichment = (Density of polyN motif at bin) / (Density of polyN motif across all bins)

Strand asymmetry bias was calculated as: (motif occurrences at non-template strand) / (motif occurrences at template strand)

The distribution of polyN motifs at the template and non-template strands relative to the TSS and the TES were calculated with bedtools *intersect* command using the gene orientation approach described earlier to generate (Figure 2c-d), (Supplementary Figure 3a-e). Bootstrapping using random sampling of genes with replacement was performed from which the standard deviation of the strand asymmetry bias was calculated.

### Template / Non-template strand asymmetries in cancer

The numbers of indels overlapping motifs found in the template or non-template strands were obtained using the bedtools *intersect* command. Enrichment was calculated for the vector of genes, reporting the number of polyN motif occurrences and the number of overlapping motifs as:

A = (indels overlapping motif at non-template) /(motif occurrences at non-template)

B = (indels overlapping motif at template) /(motif occurrences at template) Enrichment = A / (A+B) with motifs representing polyN repeat tracts of size 2-10bp and dinucleotide repeat tracts of 1-5 repeated units, at genic regions (Figure 3a-e, Figure 4a-d).

We performed bootstrapping with replacement, randomly selecting the indels overlapping motifs at template and non-template strands from each randomly-selected gene, for equal number of genes in multiple iterations, from which we calculated the standard deviation of enrichment.

MMR-deficient samples were identified using genome plots and mutational signature profiles of each patient for stomach, uterus and colorectal tumours. Subsequently, transcriptional strand asymmetry levels at indels overlapping polyT tracts were compared between MSS and MSI samples to investigate the role of mismatch repair in transcriptional strand asymmetries (Figure 3b, Supplementary Figure 6).

### RNA-seq analysis

For the comparative analysis between expression levels and transcriptional strand asymmetry, cell of origin cell lines, where available, were used from Roadmap Epigenomics project (Supplementary Table 2), (Roadmap Epigenomics Consortium et al. 2015). For each cell line, genes were grouped in expression level quantiles, namely “low”, “medium” and “high” based on the associated RPKM gene expression values.

Transcriptional strand asymmetry at indels overlapping polyN motifs was investigated in relation to gene expression levels to generate Figure 3d, Figure 4b-c, Supplementary Figure 4 c-d.

For lung cancer, using cell of origin RNA-seq data (IMR-90) from Roadmap consortium (Roadmap Epigenomics Consortium et al. 2015), polyG tracts were grouped according to their length to investigate if the length of polyG tracts was associated with transcriptional strand asymmetry at indels across the gene expression quantiles (Figure 3d).

### Replication timing analysis

Repli-Seq data for IMR-90 cell line were obtained from (The ENCODE Project Consortium, 2012) and were analysed as previously described in (Morganella et al. 2016). Genes were grouped across five replication timing quantiles and transcriptional strand asymmetry at indels overlapping polyG tracts within transcribed regions was calculated for each quantile (Figure 3e). The same type of analysis was performed for lung cancer C>A (G>T) substitutions to investigate the contribution of replication timing to the levels of transcriptional strand asymmetries at substitutions and indels overlapping polyG motifs of 2-10bp length (Figure 3e).

Statistical analyses across the manuscript were performed in python with packages “math”, “scipy”, “pandas”, “scikit-learn” and “numpy”. Figures across the manuscript were generated in python using packages “matplotlib”, “seaborn” and “pandas”.

### Author contributions

IGS, MH and SNZ conceived the concepts and analytical framework and drove the intellectual exercise. IGS, wrote the code for analysing and presenting the data. IGS, MH and SNZ wrote the manuscript with the help of GK and JJ.

## Acknowledgements

MH is supported by the Wellcome Trust Sanger Institute core grant. SNZ is funded by a CRUK Advanced Clinician Scientist Award (C60100/A23916) and a CRUK Grand Challenge Award (C60100/A25274). JJ is funded by the Swiss National Science Foundation (31003B-170267).

## References

Aken, Bronwen L., Sarah Ayling, Daniel Barrell, Laura Clarke, Valery Curwen, Susan Fairley, Julio Fernandez Banet, et al. 2016. “The Ensembl Gene Annotation System.” Database: The Journal of Biological Databases and Curation 2016 (June). https://doi.org/10.1093/database/baw093.

Baretti, Marina, and Dung T. Le. 2018. “DNA Mismatch Repair in Cancer.” Pharmacology & Therapeutics 189 (September): 45–62.

Buels, Robert, Eric Yao, Colin M. Diesh, Richard D. Hayes, Monica Munoz-Torres, Gregg Helt, David M. Goodstein, et al. 2016. “JBrowse: A Dynamic Web Platform for Genome Visualization and Analysis.” Genome Biology 17 (April): 66.

Campbell, Peter J., Gad Getz, Joshua M. Stuart, Jan O. Korbel, Lincoln D. Stein, and - ICGC/TCGA Pan-Cancer Analysis of Whole Genomes Net. 2017. “Pan-Cancer Analysis of Whole Genomes.” https://doi.org/10.1101/162784.

Delacôte, Fabien, Mingguang Han, Thomas D. Stamato, Maria Jasin, and Bernard S. Lopez. 2002. “An xrcc4 Defect or Wortmannin Stimulates Homologous Recombination Specifically Induced by Double-Strand Breaks in Mammalian Cells.” Nucleic Acids Research 30 (15): 3454–63.

Denissenko, M. F., T. B. Koudriakova, L. Smith, T. R. O’Connor, A. D. Riggs, and G. P. Pfeifer. 1998. “The p53 Codon 249 Mutational Hotspot in Hepatocellular Carcinoma Is Not Related to Selective Formation or Persistence of Aflatoxin B1 Adducts.” Oncogene 17 (23): 3007–14.

Ellegren, Hans. 2004. “Microsatellites: Simple Sequences with Complex Evolution.” Nature Reviews. Genetics 5 (6): 435–45.

Fousteri, Maria, and Leon H. F. Mullenders. 2008. “Transcription-Coupled Nucleotide Excision Repair in Mammalian Cells: Molecular Mechanisms and Biological Effects.” Cell Research 18(1): 73–84.

Francino, M. P., L. Chao, M. A. Riley, and H. Ochman. 1996. “Asymmetries Generated by Transcription-Coupled Repair in Enterobacterial Genes.” Science 272 (5258): 107–9.

Garcia-Diaz, Miguel, and Thomas A. Kunkel. 2006. “Mechanism of a Genetic Glissando: Structural Biology of Indel Mutations.” Trends in Biochemical Sciences 31 (4): 206–14.

Ghezraoui, Hind, Marion Piganeau, Benjamin Renouf, Jean-Baptiste Renaud, Annahita Sallmyr, Brian Ruis, Sehyun Oh, et al. 2014. “Chromosomal Translocations in Human Cells Are Generated by Canonical Nonhomologous End-Joining.” Molecular Cell 55 (6): 829–42.

Ginno, Paul A., Yoong Wearn Lim, Paul L. Lott, Ian Korf, and Frédéric Chédin. 2013. “GC Skew atthe 5’ and 3’ Ends of Human Genes Links R-Loop Formation to Epigenetic Regulation and Transcription Termination.” Genome Research 23 (10): 1590–1600.

Hanawalt, Philip C., and Graciela Spivak. 2008. “Transcription-Coupled DNA Repair: Two Decades of Progress and Surprises.” Nature Reviews. Molecular Cell Biology 9 (12): 958–70.

Haradhvala, Nicholas J., Paz Polak, Petar Stojanov, Kyle R. Covington, Eve Shinbrot, Julian M. Hess, Esther Rheinbay, et al. 2016. “Mutational Strand Asymmetries in Cancer Genomes Reveal Mechanisms of DNA Damage and Repair.” Cell 164 (3): 538–49.

Jasin, Maria, and Rodney Rothstein. 2013. “Repair of Strand Breaks by Homologous Recombination.” Cold Spring Harbor Perspectives in Biology 5 (11): a012740.

Kass, Elizabeth M., Pei Xin Lim, Hildur R. Helgadottir, Mary Ellen Moynahan, and Maria Jasin. 2016. “Robust Homology-Directed Repair within Mouse Mammary Tissue Is Not Specifically Affected by Brca2 Mutation.” Nature Communications 7 (1). https://doi.org/10.1038/ncomms13241.

Kunkel, Thomas A., and Dorothy A. Erie. 2015. “Eukaryotic Mismatch Repair in Relation to DNA Replication.” Annual Review of Genetics 49: 291–313.

Majewski, Jacek. 2003. “Dependence of Mutational Asymmetry on Gene-Expression Levels in the Human Genome.” American Journal of Human Genetics 73 (3): 688–92.

Morganella, Sandro, Ludmil B. Alexandrov, Dominik Glodzik, Xueqing Zou, Helen Davies, Johan Staaf, Anieta M. Sieuwerts, et al. 2016. “The Topography of Mutational Processes in Breast Cancer Genomes.” Nature Communications 7 (May): 11383.

Nik-Zainal, Serena, Ludmil B. Alexandrov, David C. Wedge, Peter Van Loo, Christopher D. Greenman, Keiran Raine, David Jones, et al. 2012. “Mutational Processes Molding the Genomes of 21 Breast Cancers.” Cell 149 (5): 979–93.

Nik-Zainal, Serena, Helen Davies, Johan Staaf, Manasa Ramakrishna, Dominik Glodzik, Xueqing Zou, Inigo Martincorena, et al. 2016. “Landscape of Somatic Mutations in 560 Breast Cancer Whole-Genome Sequences.” Nature 534 (7605): 47–54.

Petrov, Dmitri A. 2002. “Mutational Equilibrium Model of Genome Size Evolution.” Theoretical Population Biology 61 (4): 531–44.

Pleasance, Erin D., R. Keira Cheetham, Philip J. Stephens, David J. McBride, Sean J. Humphray, Chris D. Greenman, Ignacio Varela, et al. 2010. “A Comprehensive Catalogue of Somatic Mutations from a Human Cancer Genome.” Nature 463 (7278): 191–96.

Quinlan, Aaron R., and Ira M. Hall. 2010. “BEDTools: A Flexible Suite of Utilities for Comparing Genomic Features.” Bioinformatics 26 (6): 841–42.

Roadmap Epigenomics Consortium, Anshul Kundaje, Wouter Meuleman, Jason Ernst, Misha Bilenky, Angela Yen, Alireza Heravi-Moussavi, et al. 2015. “Integrative Analysis of 111 Reference Human Epigenomes.” Nature 518 (7539): 317–30.

Rodin, S., and A. Rodin. 2005. “Origins and Selection of p53 Mutations in Lung Carcinogenesis.” Seminars in Cancer Biology 15 (2): 103–12.

Romanova, Nina V., and Gray F. Crouse. 2013. “Different Roles of Eukaryotic MutS and MutL Complexes in Repair of Small Insertion and Deletion Loops in Yeast.” PLoS Genetics 9 (10):e1003920.

Sia, E. A., R. J. Kokoska, M. Dominska, P. Greenwell, and T. D. Petes. 1997. “Microsatellite Instability in Yeast: Dependence on Repeat Unit Size and DNA Mismatch Repair Genes.” Molecular and Cellular Biology 17 (5): 2851–58.

Simsek, Deniz, and Maria Jasin. 2010. “Alternative End-Joining Is Suppressed by the Canonical NHEJ Component Xrcc4–ligase IV during Chromosomal Translocation Formation.” Nature Structural & Molecular Biology 17 (4): 410–16.

Strand, M., M. C. Earley, G. F. Crouse, and T. D. Petes. 1995. “Mutations in the MSH3 Gene Preferentially Lead to Deletions within Tracts of Simple Repetitive DNA in Saccharomyces Cerevisiae.” Proceedings of the National Academy of Sciences of the United States of America 92 (22): 10418–21.

The ENCODE Project Consortium. 2012. “An Integrated Encyclopedia of DNA Elements in the Human Genome.” Nature 489 (7414): 57–74.

Tran, H. T., D. A. Gordenin, and M. A. Resnick. 1996. “The Prevention of Repeat-Associated Deletions in Saccharomyces Cerevisiae by Mismatch Repair Depends on Size and Origin of Deletions.” Genetics 143 (4): 1579–87.

Tran, H. T., J. D. Keen, M. Kricker, M. A. Resnick, and D. A. Gordenin. 1997. “Hypermutability of Homonucleotide Runs in Mismatch Repair and DNA Polymerase Proofreading Yeast Mutants.” Molecular and Cellular Biology 17 (5): 2859–65.

Zou, Xueqing, Michel Owusu, Rebecca Harris, Stephen P. Jackson, Joanna I. Loizou, and Serena Nik-Zainal. 2018. “Validating the Concept of Mutational Signatures with Isogenic Cell Models.” Nature Communications 9 (1): 1744.

