## Supplementary Material for "Transcription-coupled repair and mismatch repair contribute towards preserving genome integrity at mononucleotide repeat tracts"

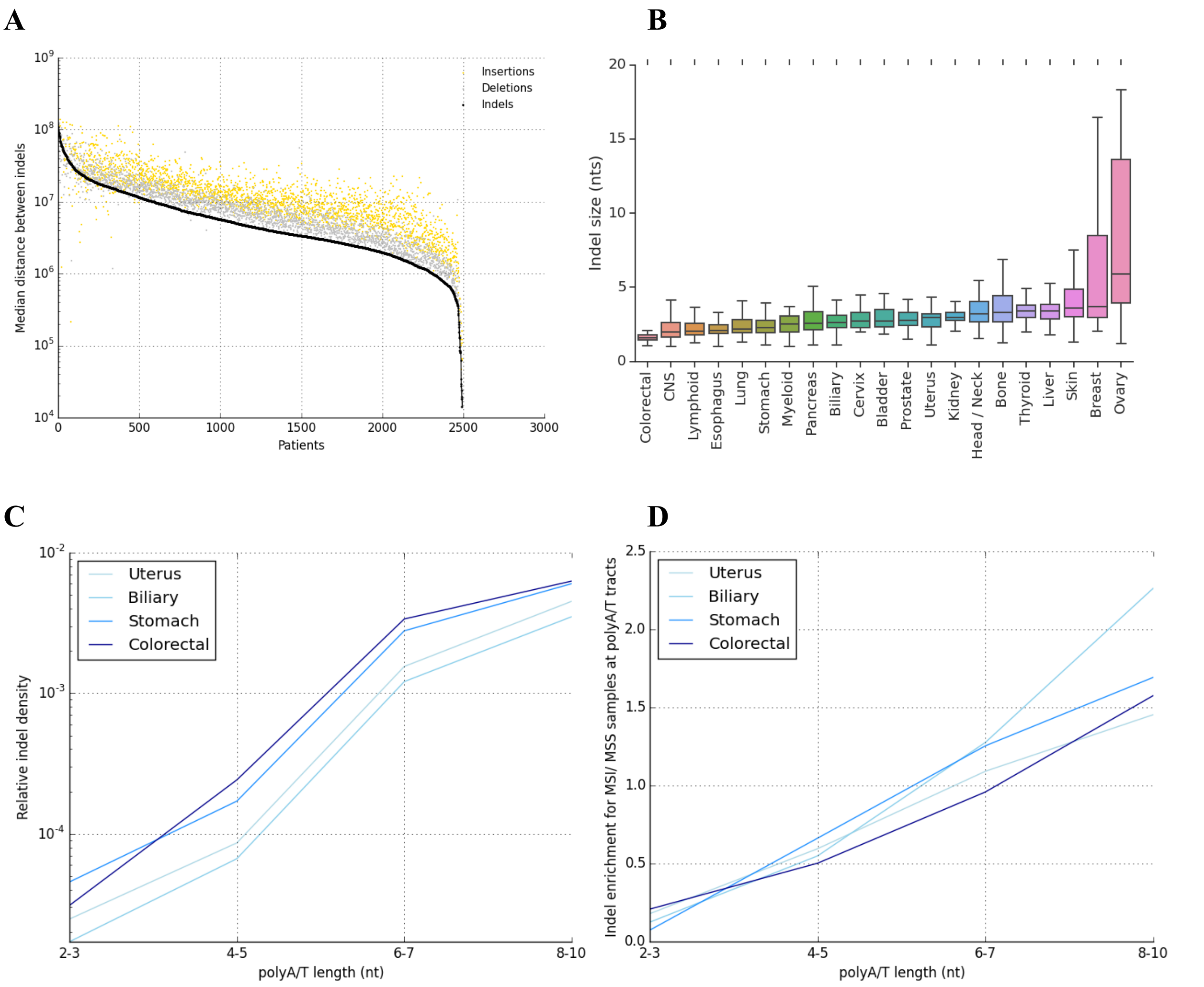


**Supplementary Figure 1: Indel size distribution patterns by patient and tumour organ. A)** Median distance between consecutive indels by patient across tumour types shown in black. Separate analysis of insertion and deletion consecutive distances shown in yellow and grey. **B)** Distribution of indel size across patients by tumour type. **C)** Relative indel density of indels at polyA/T tracts in relation to the tract length across cancer types. **D)** Mutational enrichment at MSI over MSS samples for indels at polyA/T tracts in relation to the tract length for endometrial, colorectal, biliary and stomach cancers.

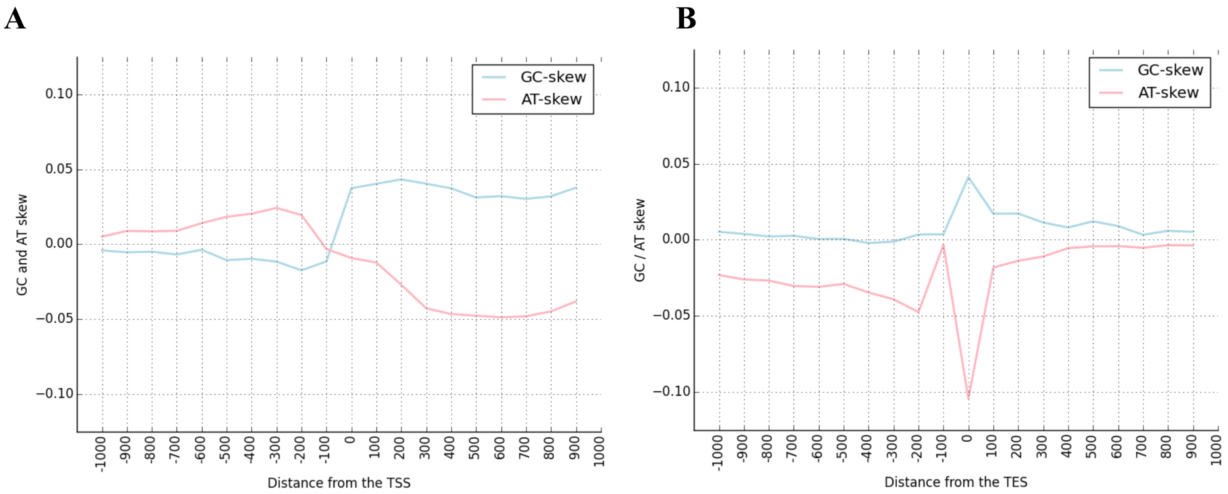


**Supplementary Figure 2: GC-skew at the TSS and TES. A)** Mean GC-skew and AT-skew around the TSS across genes, **B)** Mean GC-skew and AT-skew around the TES across genes. GC-skew defined as (G-C) / (G+C). AT-skew defined as (A-T) / (A+T).

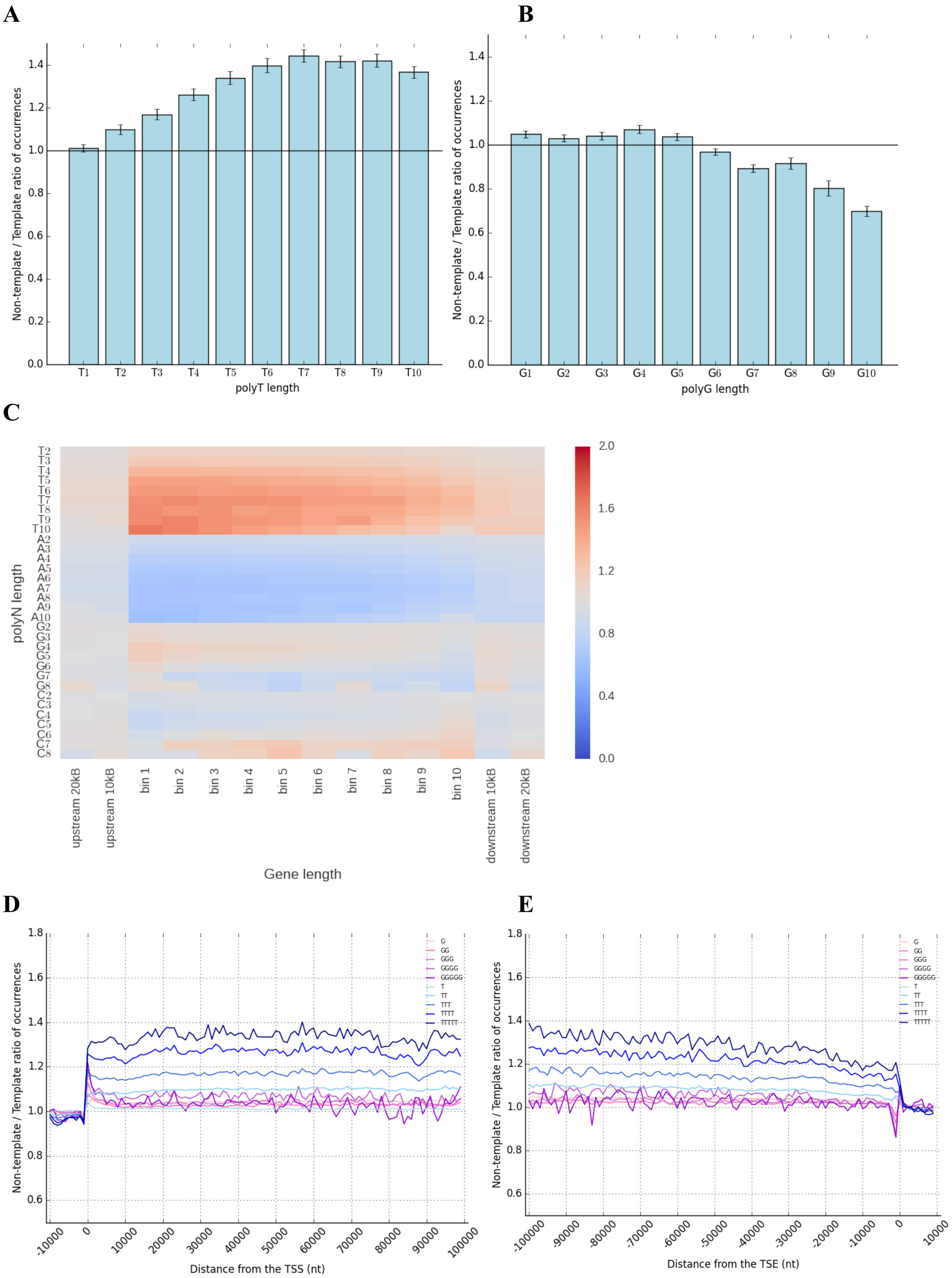
A
Non-template / Template ratio of occurrences
polyT length
T1 T2 T3 T4 T5 T6 T7 T8 T9 T10
B
Non-template / Template ratio of occurrences
polyG length
G1 G2 G3 G4 G5 G6 G7 G8 G9 G10
C
polyN length
T2 T3 T4 T5 T6 T7 T8 T9 T10 A2 A3 A4 A5 A6 A7 A8 A9 A10 G2 G3 G4 G5 G6 G7 G8 C2 C3 C4 C5 C6 C7 C8
upstream 20kB
upstream 10kB
bin 1
bin 2
bin 3
bin 4
bin 5
bin 6
bin 7
bin 8
bin 9
bin 10
downstream 10kB
downstream 20kB
Gene length
D
Non-template / Template ratio of occurrences
Distance from the TSS (nt)
G GG GGG GGGG GGGGG T TT TTT TTTT TTTTT
E
Non-template / Template ratio of occurrences
Distance from the TSE (nt)
G GG GGG GGGG GGGGG T TT TTT TTTT TTTTT

**Supplementary Figure 3: Transcriptional strand asymmetry of polyN motifs across genic regions. A)** Non-Template / Template ratio for polyT motifs within the genic region, with error bars representing 1,000-bootstrapping with replacement across genes. **B)** Non-Template / Template ratio for polyG motifs within the genic region, with error bars representing 1,000-bootstrapping with replacement across genes. **C)** Ratio of non-template to template occurrences of polyN motifs across the gene length, red indicating enrichment at non-template and blue indicating enrichment at template strand. **D)** Distance from the TSS and non-template /template polyN ratio. **E)** Distance from the TES and non-template /template polyN ratio.

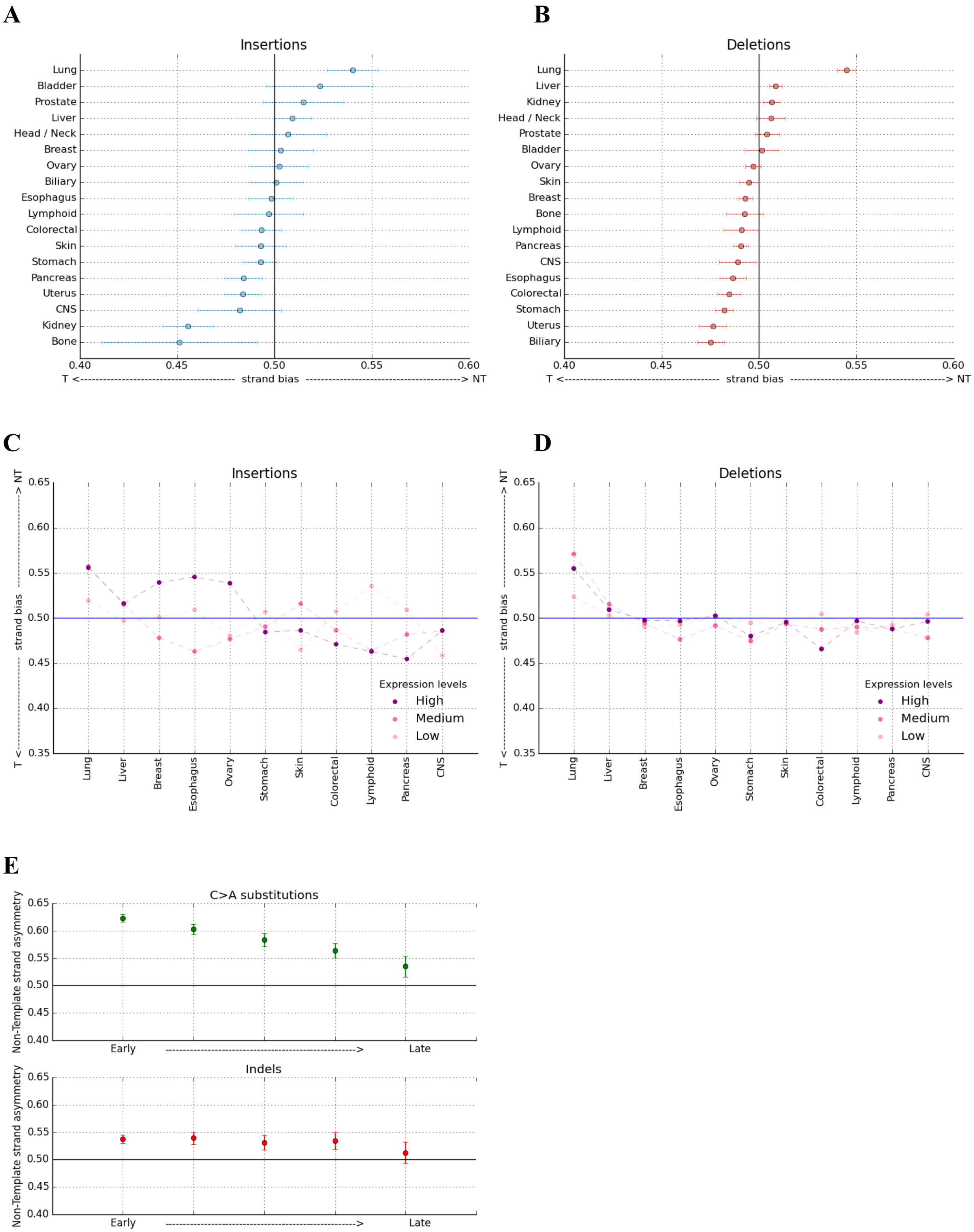


**Supplementary Figure 4: TC-NER and transcriptional strand asymmetry at indels overlapping polyG tracts. A)** Transcriptional strand asymmetry at indels overlapping polyG tracts for insertions. **B)** Transcriptional strand asymmetry at indels overlapping polyG tracts for deletions. Error bars represent standard deviation from bootstrapping with replacement. (c-d). Transcriptional strand asymmetry at indels overlapping polyG tracts across tumour organs grouped by gene expression levels for cell of origin cell lines. Transcriptional strand asymmetry at: **C)** insertions and **D)** deletions. **E)** Comparing the level of transcriptional strand asymmetry across replication timing domains for substitutions and indels in lung cancer.

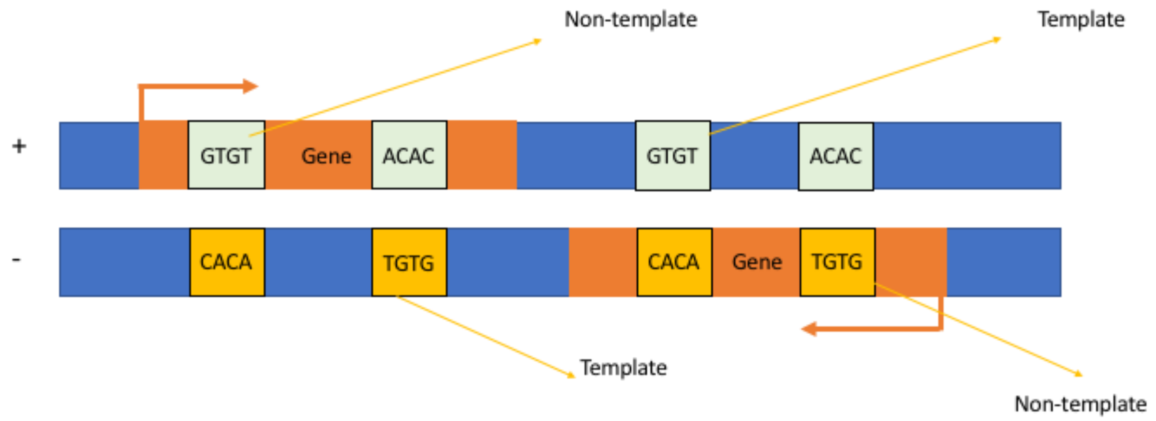


**Supplementary Figure 5: Orientation of non-overlapping dinucleotide repeat tracts relative to transcription orientation.** Example for the orientation of GTGT tracts relative to the direction of transcription for a gene on the plus strand and a gene on the minus strand, both depicted in dark orange.

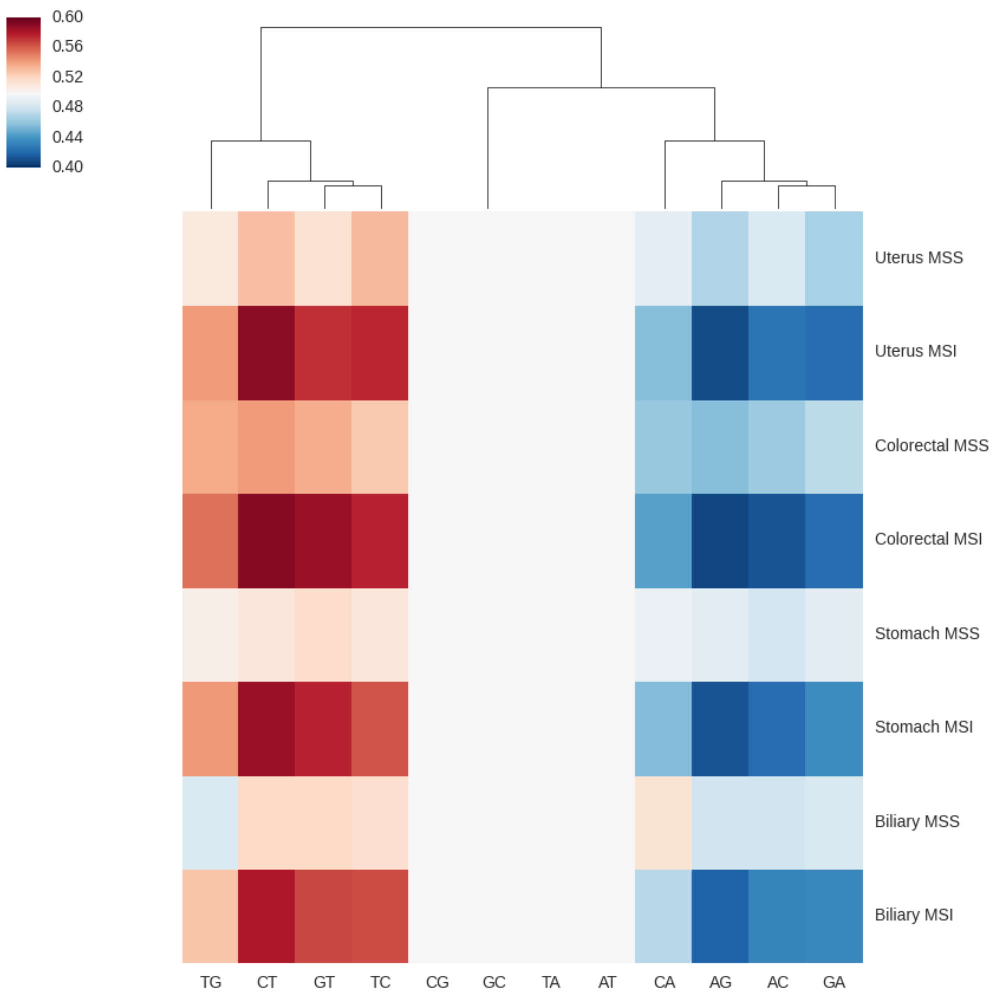


**Supplementary Figure 6: Transcriptional strand asymmetries at dinucleotide repeat motifs for MSI and MSS samples of uterus, colorectal, stomach and biliary cancers.**

A

B

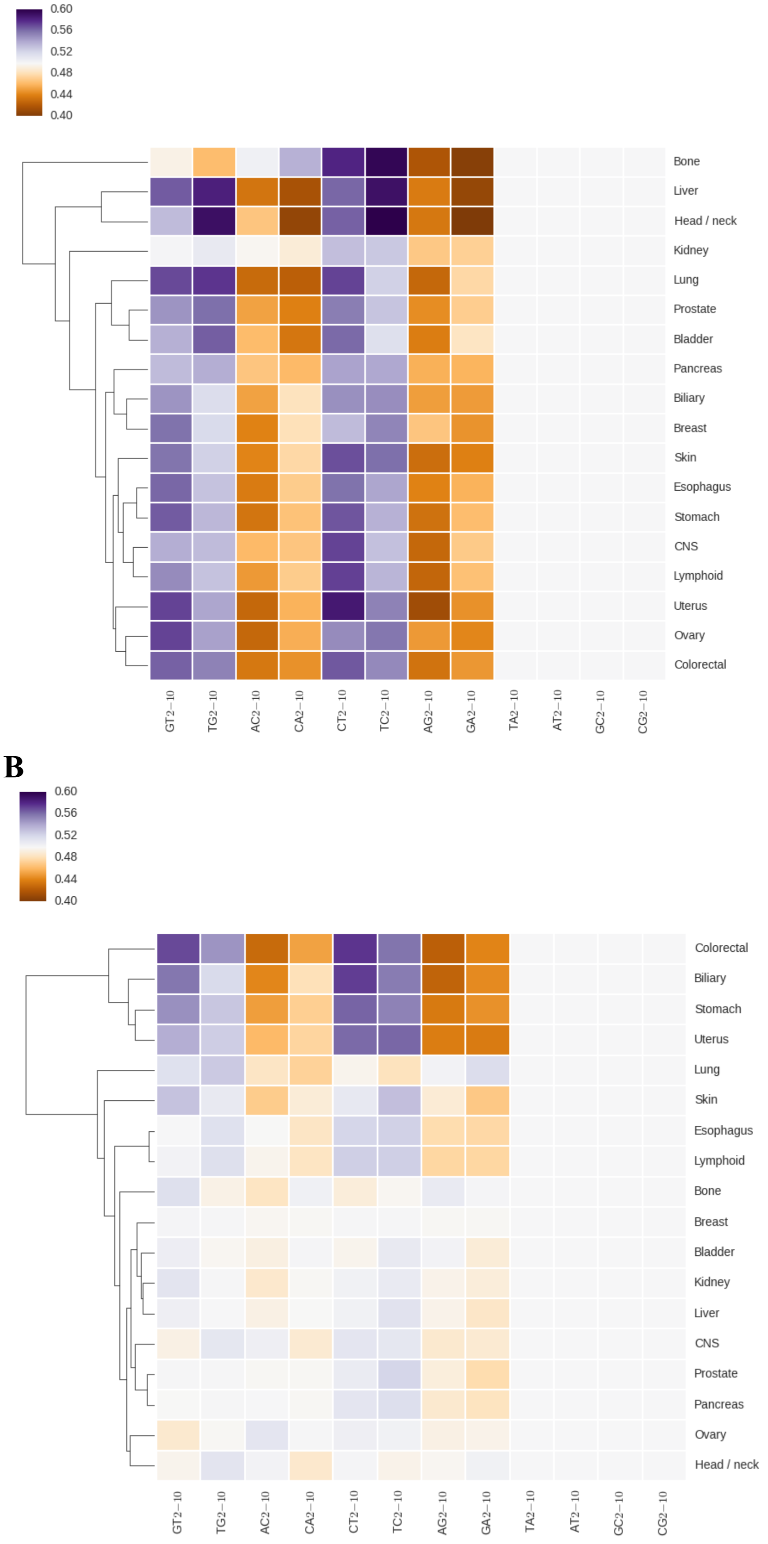


**Supplementary Figure 7: Transcriptional strand asymmetries across dinucleotide repeat motifs for A) insertions, and B) deletions across cancer types.**

**Supplementary Table 1: Number of patients, insertions and deletions per tumour organ.**

| Tumour organ | Patients | Deletions | Insertions |
|---|---|---|---|
| Bladder | 23 | 11,101 | 5,571 |
| Biliary | 34 | 119,952 | 35,024 |
| Pancreas | 313 | 93,936 | 91,392 |
| Head / Neck | 56 | 23,756 | 14,602 |
| Liver | 314 | 150,392 | 78,977 |
| Ovary | 110 | 59,917 | 27,903 |
| Prostate | 199 | 36,017 | 22,512 |
| Colorectal | 52 | 208,761 | 132,204 |
| Myeloid | 38 | 1,177 | 609 |
| Stomach | 68 | 253,355 | 62,045 |
| Cervix | 20 | 3,854 | 3,434 |
| Uterus | 44 | 119,848 | 78,578 |
| CNS | 287 | 29,362 | 19,497 |
| Lymphoid | 197 | 61,209 | 43,592 |
| Skin | 107 | 79,358 | 27,657 |
| Kidney | 186 | 104,359 | 29,518 |
| Breast | 211 | 70,333 | 23,088 |
| Esophagus | 97 | 89,741 | 63,642 |
| Thyroid | 48 | 3,101 | 1,045 |
| Bone | 89 | 14,256 | 4,527 |
| Lung | 84 | 89,842 | 34,210 |

**Supplementary Table 2:** Cell of origin RNA-seq datasets from (Roadmap Epigenomics Consortium et al. 2015) used to calculate the transcription strand asymmetry levels of cancer organs for genes of different expression levels.

| Cancer | Cell type |
|---|---|
| Breast | MCF-7 cell line |
| Colorectal | Sigmoid colon primary cells |
| Lung | IMR-90 cell line |
| Pancreas | Pancreatic primary cells |
| Liver | HepG2 cell line |
| Esophagus | Esophageal primary cells |
| Ovarian | Ovary primary cells |
| CNS | Female fetal brain cells |
| Lymphoid | K562 cell line |
| Skin | Foreskin fibroblasts |
| Stomach | Gastric primary cells |
